# Landscape structure shapes tree seedlings’ diversity at multiple spatial scales in a fragmented tropical rainforest

**DOI:** 10.1101/826339

**Authors:** Sergio Nicasio-Arzeta, Isela E. Zermeño-Hernández, Susana Maza-Villalobos, Julieta Benítez-Malvido

## Abstract

Biotic-dispersed tree seedling species are fundamental for the maintenance of the structure and function of forest patches in fragmented rainforest landscapes. Nonetheless, the effects of landscape structure and the spatial scale at which operates on seedling α- and β-diversity is unknown. Using a multi-scale approach, we assessed the relative effect of landscape composition (i.e., percentage of old-growth/secondary forest cover), configuration (i.e., aggregation/density of forest patches) and connectivity (i.e., structural and functional) on α- and β-diversity of biotic-dispersed seedlings in 16 forest patches in the Lacandona rainforest, Mexico. We assessed these effects at 13 spatial scales (from 300 to 1500 m radius, at 100 m intervals) for three α- and β-diversity orders (rare, common and dominant species). We found that patch aggregation increased species richness and reduced β-diversity of common and dominant species at similar spatial scales (500 to 600 m). Additionally, functional connectivity had a positive effect on the β-diversity of rare species in the 800 m spatial extent. These effects suggest that landscape configuration and functional connectivity sustain seedling diversity by preserving seed rain richness and the presence of large terrestrial herbivorous mammals. In contrast, the percentage of secondary forest matrix was detrimental for all α-diversity orders and the β-diversity of common and dominant species. Forthcoming conservation strategies should prevent deforestation, increase habitat amount and promote functional connectivity of forest-dependent fauna through matrix management actions.

## 1. Introduction

The long-term conservation of biodiversity is increasingly relying on novel ecosystems [1], particularly in human-modified tropical landscapes (HMTLs) [2–4]. In tropical rainforests, old-growth forest tree species are the major components of diversity, structure and function of these ecosystems [5,6], whereas the seedling community is the main regenerative pool of forest trees [7,8]. Therefore, it is not surprising that alterations of seedling species’ richness (α-diversity) and spatial dissimilarity (β-diversity) lead to the taxonomic, phylogenetic and functional impoverishment of tropical rainforests [9–11]. Accordingly, understanding how landscape structure (composition, configuration and connectivity) affects seedling diversity is crucial for biodiversity conservation in HMTLs [12,13].

A variety of studies suggest that seedling diversity is associated with the habitat amount and landscape connectivity of HMTLs [14,15]. The positive association between seeds and seedling α-diversity and the amount of old-growth forest (OGF) suggests a rescue effect of the latter by sustaining the landscape pool of colonizers and long-distance seed dispersal [16,17]. Configurational patterns that increase habitat amount, such as patch aggregation and spatial proximity (structural connectivity), can also perform a rescue effect in highly deforested landscapes [18–20].

Furthermore, in HMTLs, seedling β-diversity is sensitive to changes in abundance because of alterations in local seed dispersal and seedling recruitment [16,21,22]. In remaining forest patches, the seed rain is influenced by the surrounding secondary forests (SF), whereas seedling recruitment is sensitive to edge effects and habitat heterogeneity [9,23]. Furthermore, β-diversity is strongly affected by the functional complementarity between mammalian seed/seedling predators and seed dispersers [24,25]. Large-sized herbivorous mammals reduce the number of rare species and the abundance of dominant species at the local the scale, while dispersing the latter at the landscape scale [24,26]. The persistence of herbivorous mammals in HMTLs relies on the functional connectivity provided by a structural contrast of matrix with OGF patches [27,28]. Therefore, assessing functional connectivity is crucial in understanding the role of functional connectivity in seedling β-diversity.

Furthermore, determining the relevant scale of landscape effects’ biological responses, known as the scale of effect [29], is needed to improve conservation and management strategies. In addition, it is critical to assess the scale of effect between diversity types (α-/β-diversity) and abundance-based measurements (diversity orders) [30]. It is hypothesized that variables shaped by richness fluctuations are expected to have larger scales of effect than those affected by individuals’ abundance [31–33]. So far, only a single study has supported the above in seed rain α-diversity [16]. Thus, determining the scale of effect among diversity types and orders is required to accurately assess the response of tree seedling communities to landscape structure.

Multi-scale approaches evaluate the scale of effect of landscape structure on biological responses and the species-landscape relationships [34]. This avoids overlooking the true scale of effect if assessments were performed using very few scales within narrow ranges [29,34]. Only a handful of studies have employed multi-scale approaches for plant communities within HMTLs [16,35], and to our knowledge, there is a lack of studies on seedling communities. In this study, we employed a multi-scale approach to assess the contribution of landscape composition (OGF and SF amount), configuration (the patches’ aggregation and fragmentation) and connectivity (structural and functional) on α- and β-diversity of tree seedlings within rainforest patches. We particularly focused on the following question: Which components of landscape structure are influencing seedling α- and β-diversity and at what spatial scale?

We tested the following predictions: (1) If α-diversity is more reliant on the rescue effects of landscape structure than β-diversity, we should find a positive association between α-diversity and OGF (i.e., the amount, aggregation and structural connectivity), and a positive effect of patch fragmentation and functional connectivity for β-diversity; (2) If the SF matrix has a poor or detrimental contribution to seedlings diversity, there should be a negative effect of SF on α- and β-diversity; and (3) If the landscape structure is driving the colonization and extinction dynamics at larger spatial scales, then there should be larger scales of effect in richness-based than in abundance-based measures in α- and β-diversity.

## 2. Material and methods

### 2.1. Study site

We conducted the study at the Lacandona rainforest in southeastern Mexico (Figure 1a). The monthly temperature oscillates between 24 and 26 °C, and the annual precipitations ranges from 2500 to 3500 mm [36]. This region encompasses the largest rainforest of the Mesoamerican biodiversity hotspot [37,38]. Land-cover change to cattle pastures, however, has reduced its forest cover by more than 50% of its original extension [39,40]. We selected 16 forest patches, ranging from 1 ha to 63 ha (Figure 1b).

**Figure 1.**
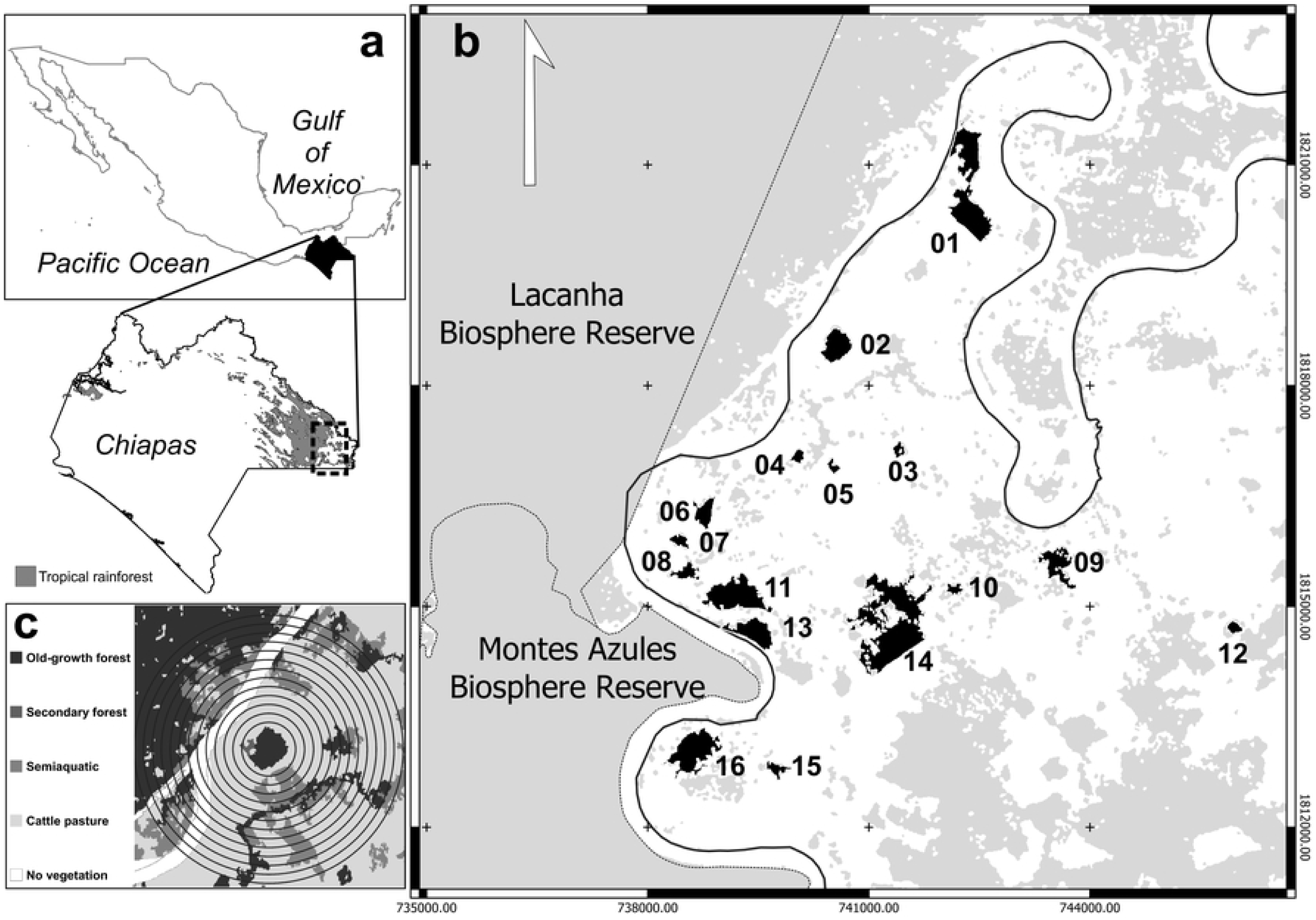
Location of the study area in the Lacandona forest in Chiapas in southeastern Mexico (a). We show the location of the 16 study forest fragments (black) in the Marqués de Comillas region (b). The 13 buffer sizes around the geographic center of the focal patch are also indicated (c).

### 2.2. Tree seedling sampling

We established ten 1-m^2^ plots arranged in a stratified random manner across a 1-ha block within each forest patch. We placed 1-m^2^ plots at least 100 m away from patch edges to avoid strong edge effects as much as possible [22,41]. Thereafter, within each plot we counted and identified all tree seedlings (10-100 cm height). We identified each seedling to the lowest possible taxonomic level with the help of a local parataxonomist and field guides [42,43]. We took samples for further identification in herbariums (MEXU, ECO-SC-H) when field identification was not possible. Furthermore, we determined the dispersal syndrome (abiotic or biotic; Table S1) of each species, based on their fruits and seed morphology [44–47]. Since in rainforests biotically-dispersed tree species comprise up to 90% of the seed rain [16,48], we developed the analysis using only this species guild. Plant nomenclature followed the Missouri Botanical Garden database Tropicos [49].

### 2.3. Diversity estimation

We assessed the sampling completeness with the sample coverage estimator of Chao and Shen [50]. We combined the data of the 10 sampling plots within each patch and estimated the proportion of the total number of individuals that belong to the species represented in the sample [51]. The sample coverage among patches was high (91.07 ± 7.47%; mean ± SD), indicating that our sampling effort was adequate for estimating species diversity [52]. We then estimated true diversity using Hill numbers [30,53]. We employed the Hill numbers of order zero (^*0*^*D*; species richness), which gives a disproportionate weight to rare species, one (^*1*^*D*; exponential Shannon index) to represent the number of typical species, and two (^*2*^*D*; inverse Simpson index) to represent the number of very abundant species [53–56]. We calculated the gamma diversity of each patch (γ_*patch*_) and the mean alpha diversity of plots (α_*plot*_) of the three diversity orders following the formula of Jost [30]. Afterward, we estimated the beta diversity between plots (^*q*^*β*_*plot*_ = ^*q*^*γ*_*patch*_/^*q*^*α*_*plot*_) to indicate the “effective number of completely distinct communities” within each patch [30]. These β-diversity values range between one (when all communities are identical) and *N* (*N* = the number of plots when all communities are completely different from each other). We used the package *vegan* [57] for the entire procedure.

### 2.4. Landscape metrics and multi-scale assessment

We employed a multispectral SPOT-5 satellite image recorded in March 2013 to carry out a supervised classification using the GRASS GIS software [58] for the entire procedure. The overall classification accuracy was 79%. We then calculated six landscape metrics (Table 1) relevant to the diversity of seed and plant communities [16,35]. The composition metrics included the percentages of old-growth forest (OGF) and secondary forest (SF) covers, whereas the configuration metrics were the density (PD) and aggregation (AI) of OGF patches. We selected the mean Euclidean-nearest neighbor distance between patches (ENN) for estimating structural connectivity and the percentage of contrasting edges (EC) as a measure of functional connectivity.[16,19,27,35,59–63]

**Table 1.** Description, metric type and ecological relevance of the landscape metrics employed in the study

| Metric | Description | Metric type | Ecological relevance | References |
| --- | --- | --- | --- | --- |
| Old-growth forest (OGF) | Percentage of landscape area covered by old-growth forest | Composition | Indicator of the landscape-scale habitat amount. It is positively associated with propagules' availability, seeds dispersal and abundance/richness of seed/seedling predators | [16,27,59–65] |
| Secondary forest (SF) | Percentage of landscape area covered by secondary forest | Composition | Measurement related to impacts on the dynamics and trajectories of floristic change within forest patches via shifts in edge effects and seed rain diversity | [9,66–69] |
| Patch density (PD) | Number of forest patches per landscape area | Configuration | Fragmentation metric positively associated with both edge effects on the tree/seedling community and with | [19,27,35,59,60,63] |
|  | (n/ha) |  | landscape connectivity for a variety of seed dispersal vectors (i.e., frugivorous bats/birds, and arboreal/terrestrial mammals) in the tropics |  |
| Aggregation index (AI) | Percentage of like-adjacencies between forest patches. Maximum aggregation indicates a single, compact patch | Configuration | A proxy of habitat amount/homogeneity in landscapes with low amounts of forest cover, influencing edge effects, local dispersal patterns and landscape complementation/supplementation dynamics | [19,66,70,71] |
| Euclidean-nearest neighbor distance (ENN) | Mean distance between forest patches within the landscape | Connectivity | Metric commonly employed to estimate structural connectivity (patch isolation). Patch isolation can have strong effects on seed dispersal success, altering tree/seedlings richness and floristic differentiation among patches | [19,20,59,61,72] |
| Edge contrast index (EC) | Average degree of edge contrast between forest patches and its immediate neighborhood | Connectivity | High percentages of contrasting edges reduce trees/seedlings survival and limit functional connectivity of seed dispersers in the tropics | [9,16,27,35,63,66,69,70,73] |

We calculated the EC using quality values that describe the permeability of matrix covers for terrestrial mammals. The permeability values were based on the assumption that overall species’ presence declines along a gradient of habitat loss and relates the percentage of each land-cover type within the landscape matrix to its relative quality [3,27]. We ranked the relative quality of each land-cover type based on the suitability of vegetation structure for mammals’ feeding, movement and/or habitat on a seven-point scale: 1 (water bodies, with the lowest suitability); 2 (anthropogenic cover); 3 (cattle pasture); 4 (arboreal crops); 5 (semiaquatic vegetation); 6 (secondary forest); and 7 (old-growth forest, representing the highest suitability). To obtain a more robust and realistic representation of landscape structure effects, we estimated the area-weighted mean of the EC index [74].

We estimated these metrics within 13 circular buffers (of a 300 to1500-m radius, at 100 m intervals) from the center of each focal forest patch (Figure 1c). The number and range of the buffers’ sizes comprise the reported scale of effect for a variety of patterns in understory vegetation, seed rain, and bird and mammal communities in HMTLs [27,35,60,62]. Although the overlapping between buffers of nearby sampling points increased at larger spatial extents, the degree of spatial autocorrelation in the model residuals is not necessarily associated with the extent of landscape overlap but rather with the proximity between sampling sites [75,76]. According to Moran’s I autocorrelation tests, we did not find spatial autocorrelation between the distance of sampling sites and the diversity patterns assessed (Table S2).

### 2.5. Statistical analyses

We assessed the scale of effect of each landscape metric using linear models. We previously verified the variables’ normality with a Shapiro-Wilk test [77]. We fitted each diversity pattern with a single landscape metric for each buffer size and assessed its predictive power with a leave-one-out cross-validation (LOOCV). Next, we calculated the proportion of the variation that can be predicted by the model using the LOOCV coefficient of determination (*R*^*2*^_*CV*_) as follows:

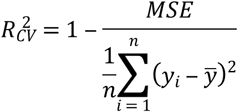

where *y*_*i*_ is the diversity value for the *i*^*th*^ patch and n is the number of patches. The *R*^*2*^_*CV*_ ranges between -∞ (indicating the model has a worse prediction power than the null model) and one (indicating the model predicts the validation data perfectly), and can be used to compare between response variables and scales of measurement [78]. We identified the scale of effect of each landscape metric by plotting the *R*^*2*^_*CV*_ of each model at each scale in function of buffer size and selected those with the strongest response [34].

We then evaluated the effects of the selected landscape metrics on diversity patterns through multiple linear models. We estimated the variance inflation factor (VIF) of landscape metrics beforehand with the *car* package [79]. Since we did not find significant collinearity (VIF ≥ 4) between the explanatory variables [80], we employed the *dredge* function of the *MuMIn* package [81] to create combinations up to three explanatory variables plus the null model (only the intercept). We ranked the models using the Akaike’s information criterion corrected for small samples (AICc) and selected those models with a AICc difference lower than two (ΔAICc < 2) as the best supported by the data [82].

Finally, we assessed the importance and the relative effect of each landscape metric measured at the scale of effect on each diversity value using an information-theoretic approach and multimodel inference [82]. For this, we selected a subset of models that had the 95% probability of containing the best model using the summed Akaike weights (*w*_*i*_) of ranked models until ∑*w*_*i*_ ≤ 0.95. We employed the *w*_*i*_ of the model’s subset to calculate the relative importance and the model-averaged parameter estimates of each explanatory variable. We carried out all statistical analysis in the R 3.5.2 statistical computing environment [83].

## 3. Results

We recorded a total of 1334 tree seedlings from 29 families, 51 genera and 72 species in 160 m^2^ (Table S3). Most seedlings were biotically-dispersed species (1258; 94.3%), belonging to 24 families, 42 genera and 58 species. Mean species density of the biotically-dispersed seedlings was 11.37 ± 2.96 species/10 m^2^ (range 6–16 species; mean ± standard error), whereas seedling density was 78.62 ± 34.03 ind/10 m^2^ (range 10–129 individuals). The most abundant species was *Inga punctata*, which represented 33 % of all individuals sampled, followed by *Ampelocera hottlei* (13%) and *Brosimum alicastrum* (11%). Less than half of the species (ca. 31%) were restricted to one patch, and only two species, *I. punctata* and *Trophis racemosa*, were found in 14 and 12 patches, respectively. Variations within α- and β-diversity orders are shown in Figure S1.

### 3.1. Scale of effect and importance of landscape structure on seedling diversity

We found α-diversity was affected by composition and structural connectivity metrics, regardless of diversity orders (Table S4). The AI and SF affected α-diversity in the 600-m buffer, whereas OGF and ENN effects occurred in smaller and larger buffer sizes, respectively (Figure S2). The AI and PD metrics affected ^1^β and ^2^β at the 500-m buffer, whereas the latter metric influenced ^0^β at the 1400 m buffer (Table S4). The EC affected ^0^β and ^2^β in the same buffer size, whereas the scale of effect of SF varied greatly between ^0^β and ^1^β (Figure S3).

When assessing the importance and contribution of the above metrics, we found that the best-fitting models of α-diversity included the positive effects of AI and the negative effects of SF (Table 2). The former metric was the second most important variable only for ^0^α (Figure 2c), whereas SF was strongly (∑*w*_*i*_ > 0.75; Figure 2a) and significantly associated with all α-diversity orders (Figure 2c). For β-diversity, we found that AI and SF metrics had negative effects for ^1^β and ^2^β, whereas EC was negatively associated with ^0^β (Table 2). The importance of these metrics was high and significant for their respective β-diversity orders (Figure 2b and 2d).

**Figure 2.**
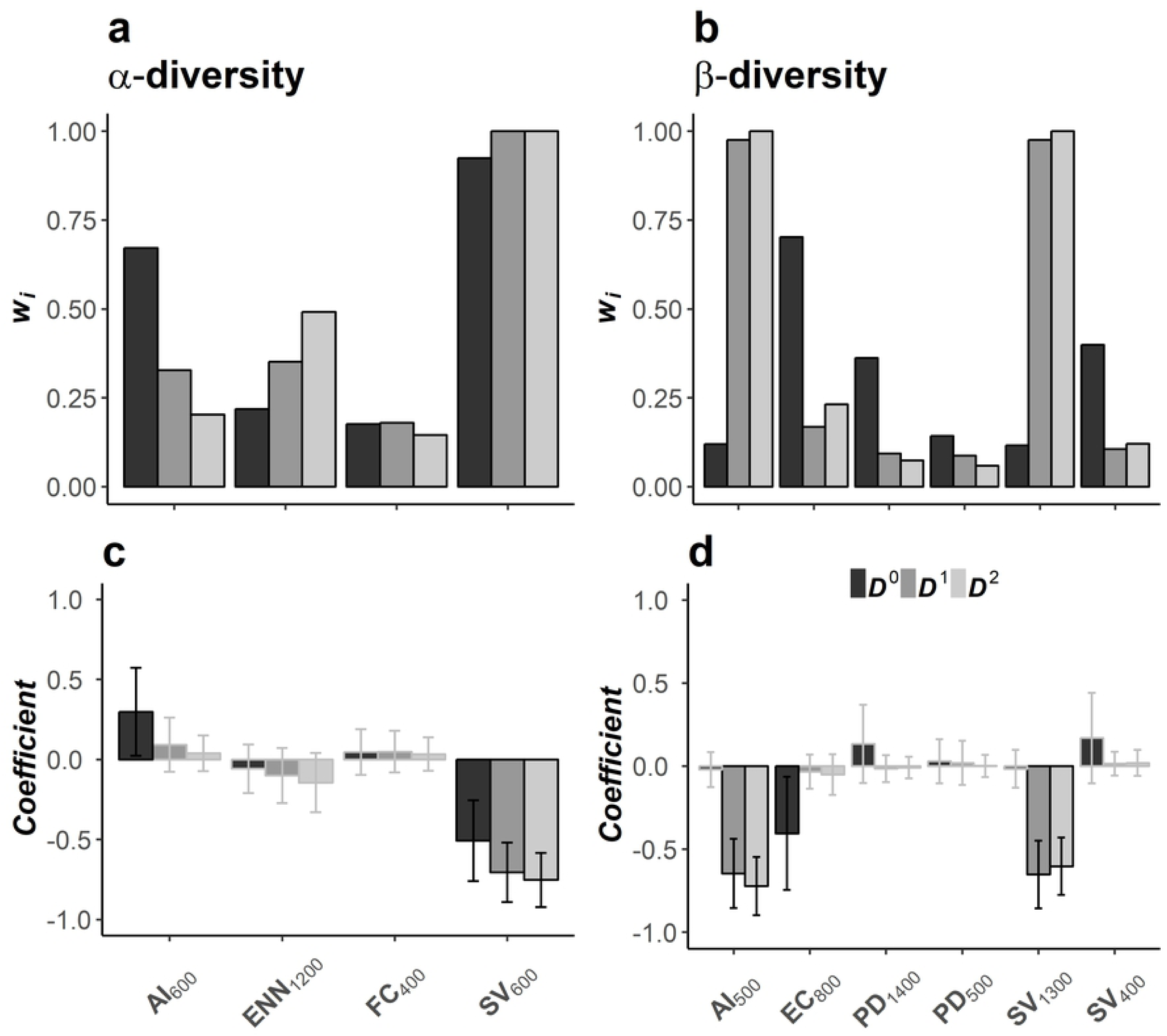
Importance and relative effects of the landscape metrics included in 95% set of models for α-(a and c) and β-diversity (b and d) in a fragmented tropical forest in southeastern Mexico. The importance of each variable is represented by the sum of the Akaike weights (∑*w*_*i*_). The effects of each covariate were estimated through a model-averaged parameter estimate of information-theoretic-based model selection and multimodel inference. The whiskers represent the unconditional standard error (USE) and the highlighted bars indicate the influential variables (those for which the USE did not include zero). The landscape metrics are the aggregation index (AI), the edge contrast index (EC), the mean distance between patches (ENN), the percentage of old-growth forest (OGF), and the percentage of secondary forest (SF). The subscript numbers indicate the scale of effect of each variable.

**Table 2.**
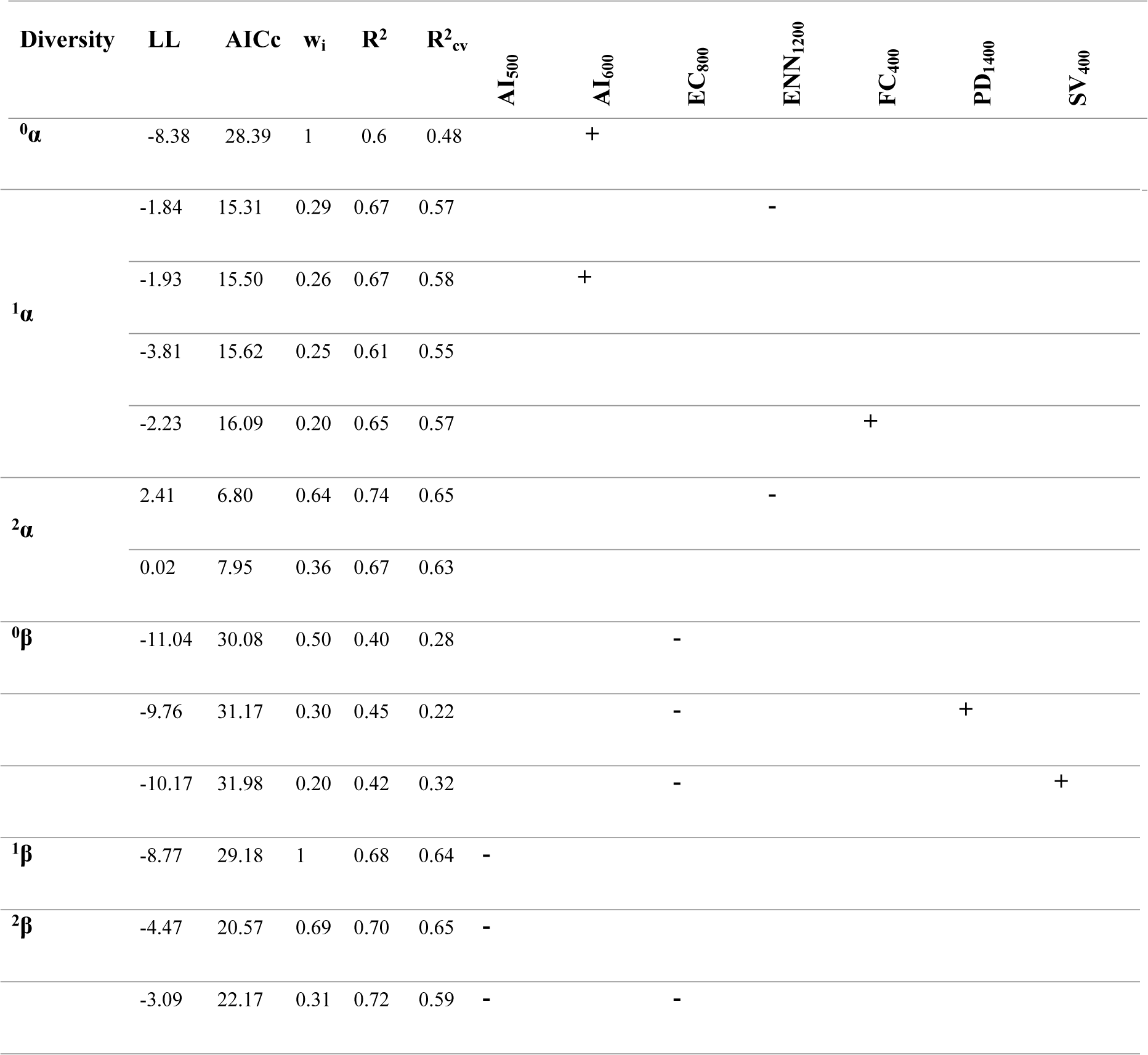
Best-supported linear models (ΔAICc < 2) that explain α- and β-diversity patterns in fragmented tropical forest in southern Mexico. The positive (+) and negative (-) symbols denote the significant effects (*P* < 0.05) of landscape metrics in the models (values reported in Table S5). The models’ log-likelihood (*LL*), the Akaike weight (*w*_*i*_), the coefficient of determination (*R*^*2*^) and the coefficient of prediction (*R*^*2*^_*CV*_) are shown. The landscape metrics are the aggregation index (AI), the edge contrast index (EC), the mean distance between patches (ENN), the percentage of old-growth forest (OGF), and the percentage of secondary forest (SF). The subscript numbers indicate the scale of effect of each variable.

## 4. Discussion

### 4.1. Landscape effects on α-diversity

As predicted, α-diversity was strongly and positively associated with habitat amount and declined as SF increased in the surrounding matrix. Nonetheless, habitat amount was related to AI rather than OGF. In addition, the AI and SF affected the three α-diversity orders in the 600-m buffer. Thus, α-diversity alterations by the above metrics and scales of effect suggest that species arrival is influenced by small-scale drivers that affect the local seed rain.

Firstly, the relationship between AI and habitat amount, as well as the observed scale of effect, is not surprising. In highly disturbed landscapes, the scale of effect is expected to be smaller since biological responses are predicted to depend on small-scale variables, such as the mean patch size [31,84]. The mean patch size is positively related to the AI in landscapes where forest cover is low (< 10%), sustaining the habitat of forest-dependent birds [18,85]. Accordingly, the AI maintains the habitat amount of seed sources and dispersers in the 600-m buffer size, where the OGF is low (24.97±10.68). This is supported by the positive effects of fragmentation within similar buffer sizes on specialist birds (564 m), primates (594-956 m) and seed rain (800 m) in HMTLs [16,60,62,86]. These species can optimize their foraging by including nearby patches within their home range as they become spatially closer [61,87]. Thus, seedling α-diversity persists through species colonization and/or recolonization from nearby patches (i.e. landscape supplementation dynamic) [70,88].

Although having non-significant effects, the scales of effect of OGF (300-500-m; Table S4) are like those observed for birds (564 m and 1261 m) and primates (200-600-m) [60,62]. Moreover, the scale of effect of a negative association between ENN and seedling richness (1200 m; Table S4) is consistent with the positive effect of OGF on seed rain richness (1244±142 m; mean ± SE) [16]. This supports the role of habitat amount in driving seedling richness through seed dispersers at smaller spatialscales and through colonization and/or extinction dynamics at larger spatialscales [16].

Secondly, the negative effect of SF on α-diversity is associated with the replacement of old-growth forest tree species by secondary forest species in the seed rain [9,23,68]. Furthermore, the richness decline of old-growth forest species is also associated to the limited movement of bird dispersers through SF within similar buffer sizes (500-564 m) [60,87]. Following seedling establishment, microclimatic alterations at forest edges promote the recruitment of disturbance-tolerant species and the mortality of shade-tolerant seedlings [22,68]. Thus, our results suggest that the secondary forest matrix reduced seedlings α-diversity through limiting seed arrival and the seedling survival of shade-tolerant species, while promoting the proliferation of disturbance-tolerant species, such as *I. punctata*.

### 4.2. Landscape effects on β-diversity

The negative association between ^1^β-/^2^β-diversity and SF and AI indicates that matrix composition and patch clumping have homogenizing effects on seedling assemblages through abundance alterations. We suggest this homogenization is driven by the input of successional-tree species at larger spatial scales and the reduction of habitat heterogeneity, seed dispersal and seedling predation at smaller scales.

The scale of effect of SF (1300-m) indicates that secondary forests are influencing tree and seedling composition, a process observed elsewhere [9,23,67,68]. In the Amazon rainforest, the abundance of successionaltree species increases in patch edges adjacent to secondary vegetation [67]. The convergence and/or divergence of tree assemblages among patches and their long-term trajectories of floristic change are driven by the extension and composition of the surrounding forest regrowth [67,68,89]. The latter are shaped by land-use history, abandonment age, forest area, and surrounding land-use types [90–92]. Our results, and the homogeneous land-use types in our study site (extensive cattle ranching), suggests that SF is converging seedling composition through the input of successional-tree species at larger scales. This supports the role of the SF as a strong driver of biotic homogenization [9,13].

Additionally, the scale of effect of the negative association between AI and ^1^β-/^2^β-diversity suggests a homogenizing effect of the habitat and inter-patch seed dispersal in the absence of terrestrial mammals. According to the “biotic differentiation hypothesis” the limited exchange of seeds and the differences of disturbance regimes across disaggregated forest patches promote a floristic differentiation in highly deforested landscapes [59,68]. The establishment of a wider array of species is facilitated due to the greater variety of conditions and a less evenly distributed number of individuals among dominant species [18,59]. Furthermore, the inverse relationship between ^0^α-diversity and ^1^β-/^2^β-diversity through AI within similar buffer sizes (600-m and 500-m, respectively), suggests the absence of terrestrial mammals. Exclusion experiments have shown that terrestrial seed and seedling predators reduce seedling ^0^α-diversity and the abundance of dominant species at the local scale [24,93]. Herbivorous mammal foraging also promotes β-diversity of dominant species through seed dispersal at larger spatial scales [24,26,94]. Terrestrial mammals’ richness is positively associated with OGF in similar buffer sizes than the observed AI effects (564 m) [27]. The above suggests an insufficient habitat amount for frugivore and browsing mammals at these spatial scales. Alongside terrestrial mammals decline, the positive effects of forest fragmentation on generalist and birds and bats sustain seedling ^0^α-diversity through local seed dispersal (Section 4.1).

Furthermore, EC affected ^0^β-diversity exclusively, supporting our assumption about the role of functional connectivity for sustaining β-diversity at local scale. This is consistent with the relationship between tree ^0^β-diversity decline and dispersal constraints of shade-tolerant species in highly fragmented landscapes [59]. In addition, the scale of effect of EC (800-m) is within the range observed for matrix quality effects on patch occupancy by terrestrial mammals in HMTLs [27,28,95]. These findings support the assumption the matrix drives functional connectivity of seed dispersers at smaller spatial scales [16]. It is noteworthy that the observed scale of effect of EC (800-m) is similar to those of matrix contrast observed for understory plants (797.88-m) and seed rain abundance (800±128 m) in other HMTLs of Mexico [16,35].

### 4.3. Conclusions and conservation implications

Our findings strengthened the evidence suggesting that conservation actions should be focused on preventing forest loss and promoting functional connectivity for maintaining forest regeneration [14,16,35]. The spatial extents for these actions should range to 500 and 800-m around patches. Accordingly, we recommend that policy interventions require preserving small OGF patches and diversifying the agricultural matrix. Implementing laws that restrict forest clearing and that promote sustainable forest management through payment for ecosystem services’ schemes can prevent further deforestation and the maintenance of remaining patches [1,96,97].

Additionally, our results also support the positive effects of fragmentation [19] for seedling β-diversity. Accordingly, promoting configurational patterns that increase habitat amount and improving matrix quality can be achieved through land-sharing schemes of smallholder agriculture [19,98]. This is supported by trend scenarios where crop intensification, the reduction of cattle ranching expansion, and the diversification of small-scale agroecosystems can reduce forest biodiversity loss in HMTLs [96,99,100].

Although our study was restricted to a single region, the study site represents a scenario of severely deforested and fragmented landscapes (less than 25% of forest cover), where the surrounding matrix is very homogeneous. Deforestation trends indicate that these landscapes will dominate HMTLs in the short term, even in the most conservative scenarios [101]. Therefore, our findings are representative of the ongoing conditions of HMTLs

Furthermore, the strong association between α-diversity with landscape composition and configuration; and β-diversity with landscape configuration and connectivity is consistent with landscape structure effects on birds and arboreal and terrestrial mammals [27,60,62,87,95]. These faunal assemblages regulate key ecological interactions, particularly seed dispersal and seed/seedling predation [24,93,102]. Therefore, the decline of forest-dependent fauna could alter seedling diversity and its long-term ecological functions, potentially compromising the ecosystem integrity of HMTLs. The above highlights the critical role of maintaining the original fauna by increasing habitat amount and functional connectivity. Nonetheless, connectivity assessments regarding these fauna assemblages are scant, and the components of matrix vegetation involved in functional connectivity are unknown. This topic represents a very important avenue for future research.

## Acknowledgments

We thank to the people of Quiringüicharo for their hospitality and support. We acknowledge the logistical and technical support provided by J. Manuel Lobato-García during the fieldwork. We thank Gilberto Jamangapé and Rafael Lombera for plant identification. Estación Chajul and Arca de Noe provided logistical assistance and accommodation. Adriana L. Luna-Nieves and Bianca A. Santini provided very valuable comments on earlier versions of the manuscript. SNA is a doctoral student from Programa de Doctorado en Ciencias Biomédicas, Universidad Nacional Autónoma de México (UNAM), and was supported by CONACyT, Mexico (scholarship 317569) and DGAPA-UNAM.

## References

1. IPBES. Global assessment report on biodiversity and ecosystem services of the Intergovernmental Science-Policy Platform on Biodiversity and Ecosystem Services. Brondizio ES, Settele J, Días S, Ngo HT, editors. Bonn, Germany: IPBES Secretariat; 2019.

2. Alroy J. Effects of habitat disturbance on tropical forest biodiversity. Proc Natl Acad Sci. 2017;114: 6056–6061. doi:10.1073/pnas.1611855114

3. Gardner TA, Barlow J, Chazdon R, Ewers RM, Harvey CA, Peres CA, et al. Prospects for tropical forest biodiversity in a human-modified world. Ecol Lett. 2009;12: 561–582. doi:10.1111/j.1461-0248.2009.01294.x

4. Kim DH, Sexton JO, Townshend JR. Accelerated deforestation in the humid tropics from the 1990s to the 2000s. Geophysical Research Letters. John Wiley & Sons, Ltd; 2015. pp. 3495–3501. doi:10.1002/2014GL062777

5. Denslow JS. Tropical Rainforest Gaps and Tree Species Diversity. Annu Rev Ecol Syst. 1987;18: 431–451. doi:10.1146/annurev.es.18.110187.002243

6. Galetti M, Bovendorp RS, Guevara R. Defaunation of large mammals leads to an increase in seed predation in the Atlantic forests. Glob Ecol Conserv. 2015;3: 824–830. doi:10.1016/j.gecco.2015.04.008

7. Webb CO, Peart DR. Seedling Density Dependence Promotes Coexistence of. Ecology. 1999;80: 2006–2017.

8. Harms KEE, Wright SJJ, Calderón O, Hernández A, Herre EA a. Pervasive density-dependent recruitment enhances seedling diversity in a tropical forest. Nature. 2000;404: 493–5.. doi:10.1038/35006630

9. Tabarelli M, Peres CA, Melo FPL. The “few winners and many losers” paradigm revisited: Emerging prospects for tropical forest biodiversity. Biol Conserv. 2012;155: 136–140. doi:10.1016/j.biocon.2012.06.020

10. Park DS, Razafindratsima OH. Anthropogenic threats can have cascading homogenizing effects on the phylogenetic and functional diversity of tropical ecosystems. Ecography (Cop). 2019;42: 148–161. doi:10.1111/ecog.03825

11. Galetti M, Dirzo R. Ecological and evolutionary consequences of living in a defaunated world. Biol Conserv. 2013;163: 1–6. doi:10.1016/j.biocon.2013.04.020

12. Perfecto I, Vandermeer J. Biodiversity conservation in tropical agroecosystems: A new conservation paradigm. Ann N Y Acad Sci. 2008;1134: 173–200. doi:10.1196/annals.1439.011

13. Arroyo-Rodríguez V, Melo FPL, Martínez-Ramos M, Bongers F, Chazdon RL, Meave JA, et al. Multiple successional pathways in human-modified tropical landscapes: new insights from forest succession, forest fragmentation and landscape ecology research. Biol Rev. 2017;92: 326–340. doi:10.1111/brv.12231

14. Santo-Silva EE, Almeida WR, Melo FPL, Zickel CS, Tabarelli M. The nature of seedling assemblages in a fragmented tropical landscape: Implications for forest regeneration. Biotropica. 2013;45: 386–394. doi:10.1111/btp.12013

15. Laurance WF, Lovejoy TE, Vasconcelos HL, Bruna EM, Didham RK, Stouffer PC, et al. Ecosystem decay of Amazonian forest fragments: A 22-year investigation. Conservation Biology. John Wiley & Sons, Ltd (10.1111); 2002. pp. 605–618. doi:10.1046/j.1523-1739.2002.01025.x

16. San-José M, Arroyo-Rodríguez V, Jordano P, Meave JA, Martínez-Ramos M. The scale of landscape effect on seed dispersal depends on both response variables and landscape predictor. Landsc Ecol. 2019; 1–12. doi:10.1007/s10980-019-00821-y

17. Charles LS, Dwyer JM, Chapman HM, Yadok BG, Mayfield MM. Landscape structure mediates zoochorous-dispersed seed rain under isolated pasture trees across distinct tropical regions. Landsc Ecol. 2019; 1–16. doi:10.1007/s10980-019-00846-3

18. Radford JQ, Bennett AF, Cheers GJ. Landscape-level thresholds of habitat cover for woodland-dependent birds. Biol Conserv. 2005;124: 317–337. doi:10.1016/j.biocon.2005.01.039

19. Fahrig L. Ecological Responses to Habitat Fragmentation Per Se. Annu Rev Ecol Evol Syst. 2017;48: annurev-ecolsys-110316-022612. doi:10.1146/annurev-ecolsys-110316-022612

20. Fahrig L. Rethinking patch size and isolation effects: The habitat amount hypothesis. Triantis K, editor. J Biogeogr. 2013;40: 1649–1663. doi:10.1111/jbi.12130

21. Benitez-Malvido J, Martinez-Ramos M. Influence of Edge Exposure on Tree Seedling Species Recruitment in Tropical Rain Forest Fragments1. Biotropica. 2003;35: 530–541. doi:10.1111/j.1744-7429.2003.tb00609.x

22. Benitez-Malvido J. Impact of forest fragmentation on seedling abundance in a tropical rain forest. Conserv Biol. 1998;12: 380–389. doi:10.1046/j.1523-1739.1998.96295.x

23. Laurance WF, Nascimento HEM, Laurance SG, Andrade AC, Fearnside PM, Ribeiro JEL, et al. Rain forest fragmentation and the proliferation of successional trees. Ecology. 2006;87: 469–482. doi:10.1890/05-0064

24. Villar N, Siqueira T, Zipparro VB, Farah F, Schmaedecke G, Hortenci L, et al. The cryptic regulation of diversity by functionally complementary large tropical forest herbivores. Edwards D, editor. J Ecol. 2019; 1365-2745.13257. doi:10.1111/1365-2745.13257

25. Kurten EL. Cascading effects of contemporaneous defaunation on tropical forest communities. Biol Conserv. 2013;163: 22–32. doi:10.1016/j.biocon.2013.04.025

26. Fragoso JM V, Silvius KM, Correa JA. Long-distance sees dispersal by tapirs increase seed survival and aggregates tropical trees. Ecology. 2003;84: 1998–2006. doi:10.1890/01-0621

27. Garmendia A, Arroyo-Rodríguez V, Estrada A, Naranjo EJ, Stoner KE. Landscape and patch attributes impacting medium- and large-sized terrestrial mammals in a fragmented rain forest. J Trop Ecol. 2013;29: 331–344. doi:10.1017/S0266467413000370

28. Cassano CR, Barlow J, Pardini R. Large Mammals in an Agroforestry Mosaic in the Brazilian Atlantic Forest. Biotropica. 2012;44: 818–825. doi:10.1111/j.1744-7429.2012.00870.x

29. Jackson HB, Fahrig L. Are ecologists conducting research at the optimal scale? Glob Ecol Biogeogr. 2015;24: 52–63. doi:10.1111/geb.12233

30. Jost L. Partitioning diversity into independent alpha and beta components. Ecology. 2007;88: 2427–2439. doi:10.1890/06-1736.1

31. Miguet P, Jackson HB, Jackson ND, Martin AE, Fahrig L. What determines the spatial extent of landscape effects on species? Landsc Ecol. 2016;31: 1177–1194. doi:10.1007/s10980-015-0314-1

32. Martin AE. The Spatial Scale of a Species’ Response to the Landscape Context Depends on which Biological Response You Measure. Curr Landsc Ecol Reports. 2018;3: 23–33. doi:10.1007/s40823-018-0030-z

33. Suárez-Castro AF, Simmonds JS, Mitchell MGE, Maron M, Rhodes JR. The Scale-Dependent Role of Biological Traits in Landscape Ecology: A Review. Curr Landsc Ecol Reports. 2018;3: 12–22. doi:10.1007/s40823-018-0031-y

34. Jackson HB, Fahrig L. What size is a biologically relevant landscape? Landsc Ecol. 2012;27: 929–941. doi:10.1007/s10980-012-9757-9

35. Hernández-Ruedas MA, Arroyo-Rodríguez V, Morante-Filho JC, Meave JA, Martínez-Ramos M. Fragmentation and matrix contrast favor understory plants through negative cascading effects on a strong competitor palm. Ecol Appl. 2018;28: 1546–1553. doi:10.1002/eap.1740

36. Pennington TD, Sarukhán J. Árboles tropicales de México. Manual para la identificación de las principales especies. 3rd ed. Mexico City: Universidad Nacional Autónoma de México; 2005.

37. González-Espinosa M, Ramírez-Marcial N, Ruiz-Montoya L. Diversidad biológica en Chiapas. Colegio de la Frontera Sur: Consejo de Ciencia y Tecnología del Estado de Chiapas, Plaza y Valdéz, editors. México; 2005.

38. Myers N, Mittermeier R a, Mittermeier CG, da Fonseca G a, Kent J. Biodiversity hotspots for conservation priorities. Nature. 2000;403: 853–8. doi:10.1038/35002501

39. Kolb M, Galicia L. Scenarios and story lines: drivers of land use change in southern Mexico. Environ Dev Sustain. 2018;20: 681–702. doi:10.1007/s10668-016-9905-5

40. Carabias J, De la Maza J, Cadena R. Conservación y desarrollo sustentable en la Selva Lacandona: 25 años de actividades y experiencia. Mexico City: Natura y Ecosistemas Mexicanos; 2015.

41. Sánchez-Gallen I, Álvarez-Sánchez FJ, Benítez-Malvido J. Structure of the advanced regeneration community in tropical rain forest fragments of Los Tuxtlas, Mexico. Biol Conserv. 2010;143: 2111–2118. doi:10.1016/j.biocon.2010.05.021

42. Martínez E, Ramos-A. CH, Chiang F. Lista florística de la Lacandona, Chiapas. Bot Sci. 1994;0: 99–177. doi:10.17129/botsci.1430

43. Sousa M. Adiciones al género Inga para la flora Mesoamericana. Acta botánica Mex. 2009;89: 25–41.

44. Ibarra-Manríquez G, Cornejo-Tenorio G. Diversidad de frutos de los árboles del bosque tropical perennifolio De México. Acta botánica Mex. 2010;104: 51–104.

45. Ibarra-Manríquez G, Martínez Ramos M, Oyama K. Seedling functional types in a lowland rain forest in Mexico. Am J Bot. 2001;88: 1801–1812. doi:10.2307/3558356

46. Durán-Fernández A, Aguirre-Rivera JR, García-Pérez J, Levy-Tacher S, De Nova-Vázquez JA. Inventario florístico de la comunidad lacandona de Nahá, Chiapas, México. Bot Sci. 2016;94: 105–121. doi:10.17129/botsci.248

47. Ibarra-Manriquez G, Oyama K. Ecological Correlates of Reproductive Traits of Mexican Rain Forest Trees. Am J Bot. 1992;79: 383. doi:10.2307/2445150

48. Jordano P. Fruits and frugivory. 2nd ed. In: Fenner M, editor. Seeds: the ecology of regeneration in plant communities. 2nd ed. Wallingford, U.K.: CABI Publishing; 2000. pp. 125–166.

49. Missouri Botanical Garden. Tropicos.org. 2019 [cited 4 Jun 2019]. Available: https://www.tropicos.org/

50. Chao A, Ma KH, Hsieh TC, Chiu C-H. User’s Guide for Online Program SpadeR (Speciesrichness Prediction And Diversity Estimation in R). Taiwan; 2016.

51. Chao A, Lee SM. Estimating the number of classes via sample coverage. J Am Stat Assoc. 1992;87: 210–217. doi:10.1080/01621459.1992.10475194

52. Chao A, Jost L. Coverage-based rarefaction and extrapolation: Standardizing samples by completeness rather than size. Ecology. 2012;93: 2533–2547. doi:10.1890/11-1952.1

53. Jost L. Entropy and diversity. Oikos. 2006;113: 363–375. doi:10.1111/j.2006.0030-1299.14714.x

54. Jost L. The relation between evenness and diversity. Diversity. 2010;2: 207–232. doi:10.3390/d2020207

55. Tuomisto H. A diversity of beta diversities: Straightening up a concept gone awry. Part 1. Defining beta diversity as a function of alpha and gamma diversity. Ecography (Cop). 2010;33: 2–22. doi:10.1111/j.1600-0587.2009.05880.x

56. Chao A, Chiu CH, Hsieh TC. Proposing a resolution to debates on diversity partitioning. Ecology. 2012;93: 2037–2051. doi:10.1890/11-1817.1

57. Oksanen J, Blanchet FG, Friendly M, Kindt R, Legendre P, McGlinn D, et al. vegan: Community Ecology Package. 2.3-3. 2019. Available: https://cran.r-project.org/web/packages/vegan/index.html

58. GRASS Development Team. Geographic Resources Analysis Support System (GRASS GIS) Software, Version 7.2. 2017. Available: http://grass.osgeo.org

59. Arroyo-Rodríguez V, Rös M, Escobar F, Melo FPL, Santos BA, Tabarelli M, et al. Plant β-diversity in fragmented rain forests: Testing floristic homogenization and differentiation hypotheses. Kitzberger T, editor. J Ecol. 2013;101: 1449–1458. doi:10.1111/1365-2745.12153

60. Carrara E, Arroyo-Rodríguez V, Vega-Rivera JH, Schondube JE, de Freitas SM, Fahrig L. Impact of landscape composition and configuration on forest specialist and generalist bird species in the fragmented Lacandona rainforest, Mexico. Biol Conserv. 2015;184: 117–126. doi:10.1016/j.biocon.2015.01.014

61. Ordóñez-Gómez JD, Arroyo-Rodríguez V, Nicasio-Arzeta S, Cristóbal-Azkarate J. Which is the appropriate scale to assess the impact of landscape spatial configuration on the diet and behavior of spider monkeys? Am J Primatol. 2015;77: 56–65. doi:10.1002/ajp.22310

62. Galán-Acedo C, Arroyo-Rodríguez V, Estrada A, Ramos-Fernández G. Drivers of the spatial scale that best predict primate responses to landscape structure. Ecography (Cop). 2018;41: 2027–2037. doi:10.1111/ecog.03632

63. Galán-Acedo C, Arroyo-Rodríguez V, Estrada A, Ramos-Fernández G. Forest cover and matrix functionality drive the abundance and reproductive success of an endangered primate in two fragmented rainforests. Landsc Ecol. 2019;34: 147–158. doi:10.1007/s10980-018-0753-6

64. Umetsu F, Paul Metzger J, Pardini R. Importance of estimating matrix quality for modeling species distribution in complex tropical landscapes: a test with Atlantic forest small mammals. Ecography (Cop). 2008;31: 359–370. doi:10.1111/j.0906-7590.2008.05302.x

65. San-José M, Arroyo-Rodríguez V, Sánchez-Cordero V. Association between small rodents and forest patch and landscape structure in the fragmented Lacandona rainforest, Mexico. Trop Conserv Sci. 2014;7: 413–432. doi:10.1177/194008291400700304

66. Harper KA, Macdonald SE, Burton PJ, Chen J, Euskirchen NIES, Brosofske KD, et al. Edge Influence on Forest Structure and Composition in Fragmented Landscapes. Conserv Biol. 2005;19: 768–782.

67. Nascimento HEM, Andrade ACS, Camargo JLC, Laurance WF, Laurance SG, Ribeiro JEL. Effects of the Surrounding Matrix on Tree Recruitment in Amazonian Forest Fragments. Conserv Biol. 2006;20: 853–860. doi:10.1111/j.1523-1739.2006.00344.x

68. Laurance WF, Nascimento HEMM, Laurance SG, Andrade A, Ewers RM, Harms KE, et al. Habitat fragmentation, variable edge effects, and the landscape-divergence hypothesis. Bennett P, editor. PLoS One. 2007;2: e1017. doi:10.1371/journal.pone.0001017

69. Ewers RM, Banks-Leite C. Fragmentation impairs the microclimate buffering effect of tropical forests. PLoS One. 2013;8: e58093. doi:10.1371/journal.pone.0058093

70. Dunning JB, Danielson BJ, Pulliam HR. Ecological processes that affect populations in complex landscapes. Oikos. 1992;65: 169–175. doi:10.2307/3544901

71. He HS, DeZonia BE, Mladenoff DJ. An aggregation index (AI) to quantify spatial patterns of landscapes. Landsc Ecol. 2000;15: 591–601. doi:10.1023/A:1008102521322

72. Rayfield B, Fortin MJ, Fall A. Connectivity for conservation: A framework to classify network measures. Ecology. 2011;92: 847–858. doi:10.1890/09-2190.1

73. Stevens SM, Husband TP. The influence of edge on small mammals: evidence from Brazilian Atlantic forest fragments. Biol Conserv. 1998;85: 1–8. doi:10.1016/S0006-3207(98)00003-2

74. Li BL, Archer S. Weighted mean patch size: A robust index for quantifying landscape structure. Ecol Modell. 1997;102: 353–361. doi:10.1016/S0304-3800(97)00071-9

75. Zuckerberg B, Desrochers A, Hochachka WM, Fink D, Koenig WD, Dickinson JL. Overlapping landscapes: A persistent, but misdirected concern when collecting and analyzing ecological data. J Wildl Manage. 2012;76: 1072–1080. doi:10.1002/jwmg.326

76. Cleary KA, Waits LP, Finegan B. Agricultural intensification alters bat assemblage composition and abundance in a dynamic Neotropical landscape. Biotropica. 2016;48: 667–676. doi:10.1111/btp.12327

77. Crawley MJ. The R Book. 2nd ed. John Wiley & Sons, Ltd.; 2013.

78. Gallardo-Cruz JA, Meave JA, González EJ, Lebrija-Trejos EE, Romero-Romero MA, Pérez-García EA, et al. Predicting tropical dry forest successional attributes from space: Is the key hidden in image texture? PLoS One. 2012;7: e30506. doi:10.1371/journal.pone.0030506

79. Fox J, Weisberg S. An R Companion to Applied Regression. 2nd ed. Thousand Oaks: SAGE Publications; 2011.

80. Kutner MH, Nachtsheim C, Neter J, Li W. Applied linear statistical models. 5th ed. McGraw-Hill; 2005.

81. Barton K. MuMIn: Multi-Model Inference. R Packag version 1421. 2018.

82. Burnham KP, Anderson DR. Model Selection and Multimodel Inference: A Practical Information-Theoretic Approach (2nd ed). Ecological Modelling. Springer-Verlag; 2002. doi:10.1016/j.ecolmodel.2003.11.004

83. R Development Core Team. R: A Language and Environment for Statistical Computing. Vienna, Austria; 2018. Available: https://www.r-project.org/

84. Ricci B, Franck P, Valantin-Morison M, Bohan DA, Lavigne C. Do species population parameters and landscape characteristics affect the relationship between local population abundance and surrounding habitat amount? Ecol Complex. 2013;15: 62–70. doi:10.1016/j.ecocom.2013.02.008

85. Bennett AF, Radford JQ, Haslem A. Properties of land mosaics: Implications for nature conservation in agricultural environments. Biol Conserv. 2006;133: 250–264. doi:10.1016/j.biocon.2006.06.008

86. Jesus FM, Pivello VR, Meirelles ST, Franco GADC, Metzger JP. The importance of landscape structure for seed dispersal in rain forest fragments. Partel M, editor. J Veg Sci. 2012;23: 1126–1136. doi:10.1111/j.1654-1103.2012.01418.x

87. Kormann UG, Hadley AS, Tscharntke T, Betts MG, Robinson WD, Scherber C. Primary rainforest amount at the landscape scale mitigates bird biodiversity loss and biotic homogenization. Maron M, editor. J Appl Ecol. 2018;55: 1288–1298. doi:10.1111/1365-2664.13084

88. Tscharntke T, Tylianakis JM, Rand TA, Didham RK, Fahrig L, Batáry P, et al. Landscape moderation of biodiversity patterns and processes - eight hypotheses. Biol Rev. 2012;87: 661–685. doi:10.1111/j.1469-185X.2011.00216.x

89. Ewers RM, Andrade A, Laurance SG, Camargo JLJL, Lovejoy TE, Laurance WF. Predicted trajectories of tree community change in Amazonian rainforest fragments. Ecography (Cop). 2017;40: 26–35. doi:10.1111/ecog.02585

90. Zermeño-Hernández I, Méndez-Toribio M, Siebe C, Benítez-Malvido J, Martínez-Ramos M. Ecological disturbance regimes caused by agricultural land uses and their effects on tropical forest regeneration. Hölzel N, editor. Appl Veg Sci. 2015;18: 443–455. doi:10.1111/avsc.12161

91. Zermeño-Hernández I, Pingarroni A, Martínez-Ramos M. Agricultural land-use diversity and forest regeneration potential in human- modified tropical landscapes. Agric Ecosyst Environ. 2016;230: 210–220. doi:10.1016/j.agee.2016.06.007

92. Chazdon RL. Tropical forest recovery: Legacies of human impact and natural disturbances. Perspect Plant Ecol Evol Syst. 2003;6: 51–71. doi:10.1078/1433-8319-00042

93. Camargo-Sanabria AA, Mendoza E, Guevara R, Martínez-Rmos M, Dirzo R. Experimental defaunation of terrestrial mammalian herbivores alters tropical rainforest understorey diversity. Proc R Soc B Biol Sci. 2014;282. doi:10.1098/rspb.2014.2580

94. Fragoso JM V., Huffman JM. Seed-dispersal and seedling recruitment patterns by the last Neotropical megafaunal element in Amazonia, the tapir. J Trop Ecol. 2000;16: 369–385. doi:10.1017/S0266467400001462

95. Thornton DH, Branch LC, Sunquist ME. The relative influence of habitat loss and fragmentation: Do tropical mammals meet the temperate paradigm? Ecol Appl. 2011;21: 2324–2333. doi:10.1890/10-2124.1

96. DeClerck FAJ, Chazdon R, Holl KD, Milder JC, Finegan B, Martinez-Salinas A, et al. Biodiversity conservation in human-modified landscapes of Mesoamerica: Past, present and future. Biol Conserv. 2010;143: 2301–2313. doi:10.1016/j.biocon.2010.03.026

97. Ricketts TH, Daily GC, Ehrlich PR, Michener CD. Economic value of tropical forest to coffee production. Proc Natl Acad Sci U S A. 2004;101: 12579–12582. doi:10.1073/pnas.0405147101

98. Pazos-Almada B, Bray DB. Community-based land sparing: Territorial land-use zoning and forest management in the Sierra Norte of Oaxaca, Mexico. Land use policy. 2018;78: 219–226. doi:10.1016/j.landusepol.2018.06.056

99. Chazdon RL, Harvey CA, Komar O, Griffith DM, Ferguson BG, Martínez-Ramos M, et al. Beyond reserves: A research agenda for conserving biodiversity in human-modified tropical landscapes. Biotropica. 2009;41: 142–153. doi:10.1111/j.1744-7429.2008.00471.x

100. IPBES. The Regional Assessment Report on Biodiversity and Ecosystem Services for the Americas. Rice J, Seixas CS, Zaccagini ME, Bedoya-Gaitán M, Valderrama N, editors. Bonn, Germany; 2018.

101. Taubert F, Fischer R, Groeneveld J, Lehmann S, Müller MS, Rödig E, et al. Global patterns of tropical forest fragmentation. Nature. 2018;554: 519–522. doi:10.1038/nature25508

102. Breitbach N, Laube I, Steffan-Dewenter I, Böhning-Gaese K. Bird diversity and seed dispersal along a human land-use gradient: high seed removal in structurally simple farmland. Oecologia. 2010;162: 965–76. doi:10.1007/s00442-009-1547-y

